# Population networks from DNA sequences: methodological developments

**DOI:** 10.1101/188268

**Authors:** B. Schaeffer, V. Nicolas, F. Austerlitz, C. Larédo

## Abstract

Several classes of methods have been proposed for inferring the history of populations from genetic polymorphism data. As connectivity is a key factor to explain the structure of populations, several graph-based methods have been developed to this aim, using population genetics data. Here we propose an original method based on graphical models that uses DNA sequences to provide relationships between populations. We tested our method on various simulated data sets, describing typical demographic scenarios, for different parameters values. We found that our method behaved noticeably well for realistic demographic evolutionary processes and recovered suitably the migration processes. Our method provides thus a complementary tool for investigating population history based on genetic material.

## 1 Introduction

In population genetics, a central challenge is to determine the structure and history of populations and, to this aim, various methods have been proposed. In particular, several graph-based methods were developed to find structure among populations and patterns of gene flow. For example, Dyer and Nason [2004] used the notion of conditional independence, Rozenfeld et al. [2008] proposed a method based on the search of the percolation point and Jombart et al. [2008] added a spatial component to a principal component analysis. Many authors also designed computer programs simulating complex demographic histories (e.g. [Laval and Exco ffier, 2004], [Guillot, 2009]). Besides coalescent-based methods [Kingman, 1982] were developed to infer the history of populations. For instance, the software Ima2 [Hey, 2010] can infer the history of separation and migration of a set of populations. Nevertheless, as these methods are highly computer intensive, they cannot typically handle more than ten populations, while phylogeographic studies rely often on a much larger set of populations.

In this paper, we considered a data set composed of populations for which DNA sequences had been collected. Our aim was to investigate potential patterns of relationships between these populations. A means to find out these relationships is to use random graphs and network analysis. A network consists of a graph where each node represents an entity (here a well identified population characterized by some DNA sequences) and where edges between two nodes correspond to any relationship between these entities (populations). Using the Graphical Models framework (e.g. [Whittaker, 1990], [Lauritzen, 1996]), we developed a statistical method for inferring relationships among populations from DNA sequences. It is based on conditional independence, which explore whether a relationship between two populations exists taking into account all the other populations. Using conditional independence for population graphs was first proposed by Dyer and Nason [2004] for allozyme data. From these data, the authors used an encoding stage, PCA technics and different steps of normalization to get a correlation matrix. We propose here a generic and original method which allows getting the correlation matrix directly from the genetic sequences. Another major difference lies in the statistical testing procedure required to determine which elements of the partial correlation matrix are significantly greater than zero. Dyer and Nason [2004] used the edge exclusion deviance (EED) technique whereas we chose to use the Fisher Z-transformation which turns the sampling distribution of correlation coe cients into an approximate normal distribution. We assessed the performance of our method on simulated data.

The paper is organized as follows. Section 2 contains a short recap on Graphical Models. Section 3 describes precisely how we build populations networks from DNA sequences. Section 4 is devoted to simulations of various evolutionary population models which were analysed using our method. Section 5 contains the discussion of the results on simulations data according to the characteristics of the evolutionary scenarios (effective population sizes, mutation rates, times since the MRCA, separation times, migration rates).

## 2 Short recap on graphical models

A graphical model is a family of joint probability distributions for multivariate random observations defined in terms of conditional independences and represented by a graph (see Whittaker [1990], Lauritzen [1996]). A graph *𝒢* = (*𝒦*, ε) is composed of a finite set *𝒦* of nodes with cardinality *K* and a set of edges Ε which describes the links between pairs of distinct nodes. A random variable *X_υ_* is associated to each node υ, and we denote by (*X_υ_*,υ ∈ *𝒦*) the random vector composed of these *K* random variables. The edges of the graph can be directed or undirected. Here, we considered undirected edges which correspond thus to an undirected graph. A graph can be represented by a *K × K* adjacency matrix which takes values in {0, 1}. The entry at (*k*, *k*′)_{*k*≠*k*′}_ is {1} if there is an edge between nodes *k* and *k*′; Considering the joint probability distribution of the random variables (*X_*k*_,*k* ∈ *𝒦**), the presence of an edge between two nodes *k*, *k*′ implies a conditional dependency between the two variables *X_*k*_, X_*k*′_*; the absence of an edge between *k*, *k*′ occurs if the variables *X_*k*_, X_*k*′_* are independent given all the remaining variables. This property called “pairwise Markov Property” defines an independence graph.

In Gaussian Graphical Models (GGM), the random variable (*X_υ_*,υ ∈ *𝒦*) has a multivariate Normal distribution. The pairwise Markov property of two nodes *k*, *k*′ simplifies into a null partial covariance or correlation between the random variables *X_*k*_ and X_*k*′_* conditionally on all the remaining random variables.

Given that *𝒦* has cardinality *K*, let us denote by Σ = (Σ_*kk*′_) the *K* × *K* covariance matrix of the Gaussian random variable (*X_υ_*,υ ∈ *𝒦*) and by *R* = (ρ_*kk*′_)_1≤*k*,*k*′≤*K*_ its correlation matrix. Actually, the partial correlation matrix Π = (π_*kk*′_)_1≤*k*,*k*′≤*K*_ has a simple explicit expression based on Ω = (ω_*kk*′_)_1≤*k*,*k*′≤*K*_, the inverse of the correlation matrix *R*,

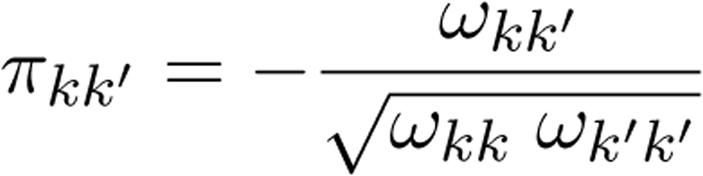

Inferring an independence graph consists in detecting the non-null partial correlation coefficients between pairs of random variables (*X_υ_*,*υ* ∈ *𝒦*) using multiple testing procedures based on an *L*-sample of *X*. For each pair of nodes *k*, *k*′, we test the null hypothesis *H_0_*(*k*, *k*′) : “there is conditional independency between the two random variables *X_k_, X_k′_*” (implying π_*kk*′_ = 0) against the alternative *H_1_*(*k*, *k*′): “there is no conditional independency between *X_*k*_, X_*k*′_*”. Rejecting *H*_0_(*k*, *k*′) means that the two variables *X_*k*_, X_*k*′_* interact in some way. In that case, we set to 1 the *kk*′-element of the adjacency matrix and draw an edge between the two nodes *k* and *k*′; otherwise we set this element to 0 and no edge is drawn. Therefore, the *K*(*K* − 1)/2 tests based on the values π_*kk*′_ of the partial correlation matrix have to be performed in order to infer the adjacency matrix.

To test the correlation coe cients we chose to use the Fisher Z-transform *u* ∈ [0,1] → 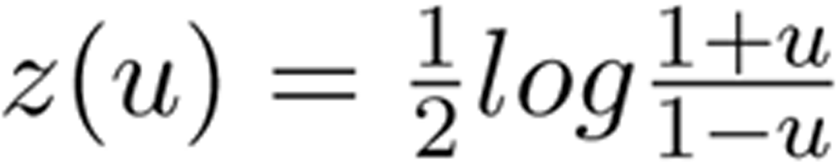 for stabilizing the variance [Fisher, 1924]. Indeed, if (*X, Y*) is a two dimensional Gaussian random variable such that *corr(X, Y) = ρ*, then the Pearson correlation estimator *r* of ρ built on an *L*-sample *(X_l_, Y_l_)_1≤l≤L_* of (*X, Y*) satisfies that *z(r_*kk*′_)* is approximately Gaussian with mean 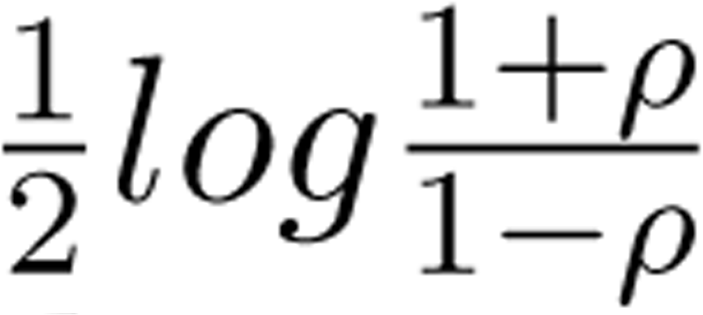 and variance 1/(*L* − 3). Similarly, if *X* is a *K-* dimensional Gaussian random variable with partial correlation coe cients π_*kk*′_, the estimator 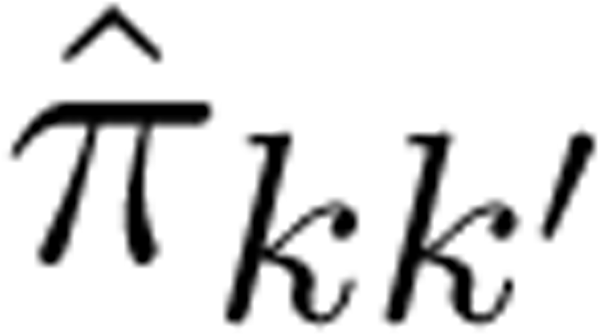 of π_*kk*′_ built on an *L*-sample of *X* satisfies that *z*(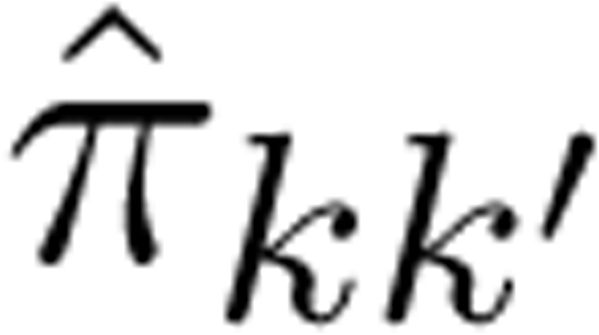) is approximately Gaussian with mean *z(π_*kk*′_)* and variance 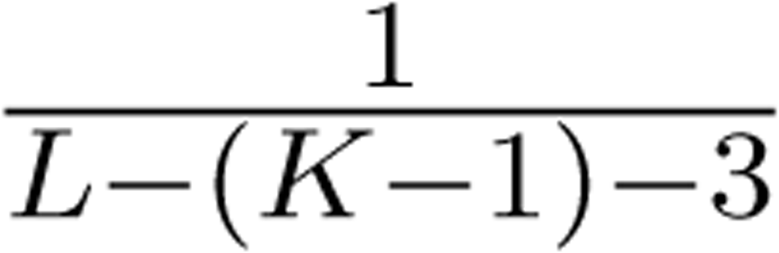 (see [Anderson, 2003]). Therefore, choosing the Fisher *Z*-transform to test the partial correlation coe cients stabilizes the variance and thus leads to better testing procedures.

For each pair of nodes *k*, *k*′, an estimator 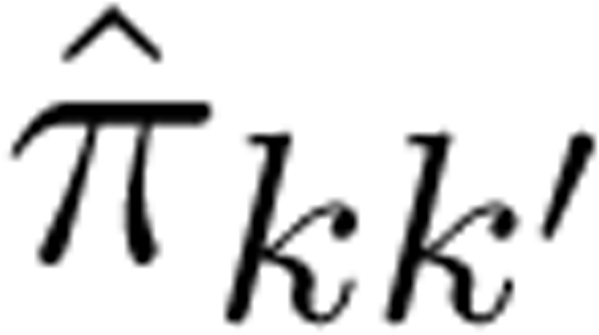 of π_*kk*′_ is computed together with a *p*-value under the null hypothesis; *H*_0_(*k*, *k*′) is rejected if this *p*-value is smaller than a prescribed threshold α_π_. The graph is inferred from the results of all these 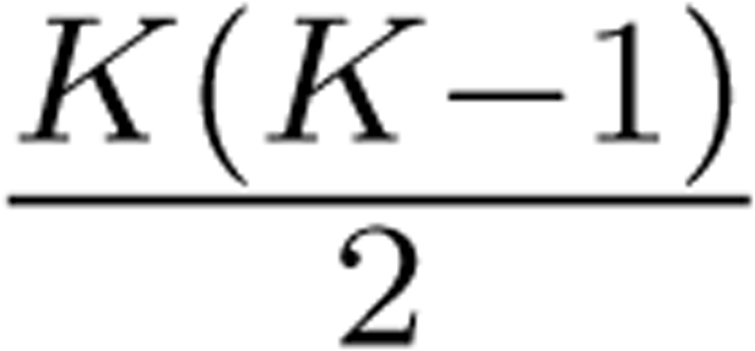 tests. In order to keep the significance level of the independence graph to a given level α, the *p*-values computed for all these tests have to be adjusted. In this study, we used the Bonferroni correction [Bonferroni, 1937] which is a strong conservative method to control the probability of false discoveries when performing multiple tests. The False Discovery Rate procedures [Benjamini and Hochberg, 1995], mostly used for multiple hypothesis testing, are less appropriate in this framework because they require independence between tests which does not hold for graphs [Korn et al., 2004].

## 3 Building a population network from DNA sequence data

We consider a set of populations comprising N specimens for which one or more genes have been sequenced. These specimens are partitioned into the *K* well identified populations. Our aim is to find potential links between these populations using the graphical model statistical framework. The method developed in this paper is a non parametric statistical method: it is not based on genetic population model, but relies on the observed DNA sequences and their variability (which clearly derives from many phenomena).

### 3.1 Data cleansing

Only the polymorphic sites were considered since they are the only informative sites. Here, we considered them as independent random variables providing information on the relationships between the populations. In what follows, let *k* (*k* = 1,…, *K*) denotes the index of a population, *L* the number of polymorphic sites and *i*(*i* = 1,…, *L*) the *i*-th polymorphic site. Each nucleotide *A* (resp. *C, G, T*) is described by an index *r* = 1 (resp. 2, 3, 4).

### 3.2 Distances between populations

Each population *k* can be described by a random vector *X*_*k*_ = (*X*_*k*_(*i,r*)) of dimension 4L ({1,…, *L*} × 1,…, 4) with values in [0,1]^4*L*^ having the probability distribution *Q*_*k*_ = *P* (*X*_*k*_ = *x*_*k*_) = *P* (*X*_*k*_(*i, r*) = *x*_*k*_(*i, r*), 1 ≤ *i* ≤ *L*, 1 ≤ *r* ≤ 4). This distribution characterizes a population.

From the data set, we can consider an estimate of this distribution, the natural one being the empirical estimate obtained as follows. First, at a polymorphic site *i*, compute the vector containing the allele frequencies (*pk*,*i*,1, *pk*,*i*,2, *pk*,*i*,3, *pk*,*i*,4) of A, C, G, T, then estimate *Q*_*k*_ by

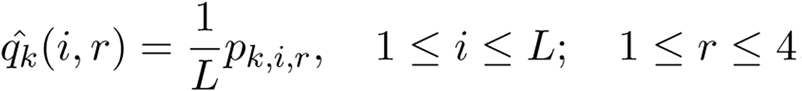

The distance between two populations *k*, *k*′ is measured by means of the Hellinger distance *H*^2^ between the two probability distributions *Qk* and *Qk*′ defined as

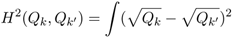

Indeed, given two probability distributions, the Hellinger distance is widely used in statistics to measure their difference. This distance is especially well adapted for hypothesis testing and is moreover convenient for considering product measures. An estimate of it for two populations is

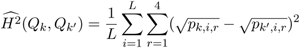

Up to a constant, this distance is equal to the genetic distance proposed by Cavalli-Sforza and Edwards [1967] between populations. It is also the Euclidean distance between 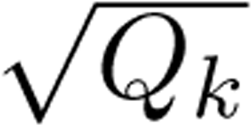 and 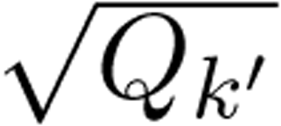 considered as vectors in *R*^4*L*^ and therefore, inherits its properties. The Hellinger distance is often expressed using the Hellinger a nity between two probabilities 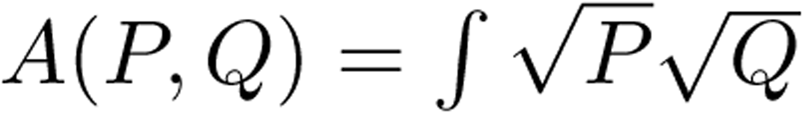 since *H*^2^(*P, Q*) = 2 − 2*A*(*P, Q*). Here *A*(Q_*k*_, Q_*k*′_) can be written as

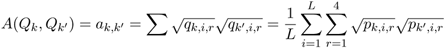

The quantity *a*_*k*,*k*′_ is similar to an association coe cient between two populations *k* and *k*′. It takes values in the interval [0, 1], reaches its maximum value {1} if the two distributions are identical, and satisfies

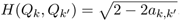

### 3.3 Correlations between populations

Taking into account the specific structure of the distributions *Q*_*k*_, *k* = 1,…, *K* along the sites, we defined the empirical covariances 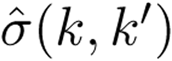 and correlations 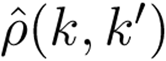 between two populations *k* and *k*′ as follows. Let first define

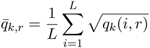

Then the empirical covariance and correlation coefficients are

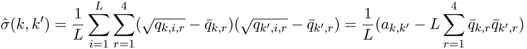

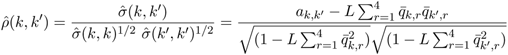

This is similar to Gower [1966] who derives the association coefficient and the correlation between two points from a distance d defined on an arbitrary space.

### 3.4 Relationships between populations

To infer relationships (edges) between populations, we have to detect conditional dependencies. We used the GGM formalism described above. From the empirical correlation matrix *R* = (ρ_*kk*′_), we calculated the partial correlation matrix Π = (ρ_*kk*′_) and tested each element of this matrix. Tests were performed using the Fisher’s Z-transformation of the partial correlation: 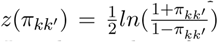 ([Fisher, 1924]). Its approximate variance is 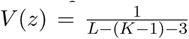 where *L* is the number of sites and *K* the number of populations. Then, the *p*-values under the null hypothesis were calculated and adjusted with the Bonferroni correction [Bonferroni, 1937]. The null hypothesis is rejected when the adjusted-*p*-value is smaller than α. An adjacency matrix is obtained accordingly, and a graph is drawn.

## 4 Simulations

### 4.1 Design of simulations

Using the SIMCOAL2 program [Laval and Excoffier, 2004], five populations (indexed by *k*, *k* = {1, …, 5}) with effective population size *N* = 1 000 were simulated under several symmetric migration models: a model with no migration, an Island model [Wright, 1931], a linear Stepping Stone model [Kimura and Weiss, 1964] and an expansion model. For these four models, the populations split simultaneously *T* generations ago. In the Islandmodel, migrations occurred symetrically between all pairs of populations. In the linear Stepping Stone model, migrations occurred symetrically between populations *k* and *k* + 1, *k* ∈ {1, 2, 3, 4, 5}. In the expansion model, some migrants from population 1 have generated 4 new populations, then migrations occured only from population 1 to each other. A fifth model called the multi-T model was tested. In this last model, the populations split at various times, *T*, *T*/2, *T*/5 and *T*/10, without migration. In all simulations, we set the length of DNA sequences to *L* = 1 000 sites and the sample sizes to *s* = 25. We varied splitting times *T*, *T* ∈ {100 000, 1 000 000} and mutation rates μ, μ ∈ {2.5 × 10^−6^, 12.5 × 10^−6^, 2.5 × 10^−5^} with a transition/transversion ratio of 9/1, assuming a finite-site mutation model. These mutation rates correspond to intraspecific thetas (θ = 2*N*μ) equal to 5, 25 and 50 respectively. In the Island, linear Stepping Stone and expansion models, two migration rates m were tested, *m* = {0.0005, 0.005}.

For each evolutionary scenario, we performed 1 000 simulations. Each simulation yielded a sequence data set. We applied our method on each data set, yielding a graph. In all cases α, was set to 0.05. The results are represented in figures by graphs and histograms. On graphs, solid lines correspond to edges detected in more than 80% of the simulations and dashed lines correspond to edges detected between 60 and 80% of the simulations. The number of solid and dashed lines in the graphs can be found in the histograms. For example in Figure 1, let us consider the histogram and the graph corresponding to the linear Stepping Stone model for θ = 5; the last bar of the histogram lies from 0.9 to 1.0 on the x-axis and reaches 2 on the y-axis. This means that two links were detected in more than 80% of the simulations. These two links are represented with solid lines in the graph above. On the same histogram, another bar lies from 0.7 to 0.8 on the x-axis and reaches 2 on the y-axis. This means that two links were detected between 60 and 80% of the simulations. These two links are represented with dashed lines in the graph above. All other links were detected in less than 60% of the simulations. So no link was added to this graph.

### 4.2 Simulation results

Figure 1 (resp. 2) corresponds to *T* = 1 000 000 (resp. *T* = 100 000), *s* = 25 and *m* = 0.0005 (for models with migration). For the three models with migration, results were similar for *T* = 100 000 and *T* = 1 000 000. While no edges were detected in the Island model when θ = 5, the occurrence probability of edges increased with θ (see the histograms in the figures). For the linear Stepping Stone and expansion models, the graphs reflected quite well the migration patterns. However for the linear Stepping Stone model and small θ, the detection probabilities of edges 2-3 and 3-4 ranged from 0.7 to 0.8. Similar results were obtained for *m* = 0.005 (not shown).

**Figure 1:**
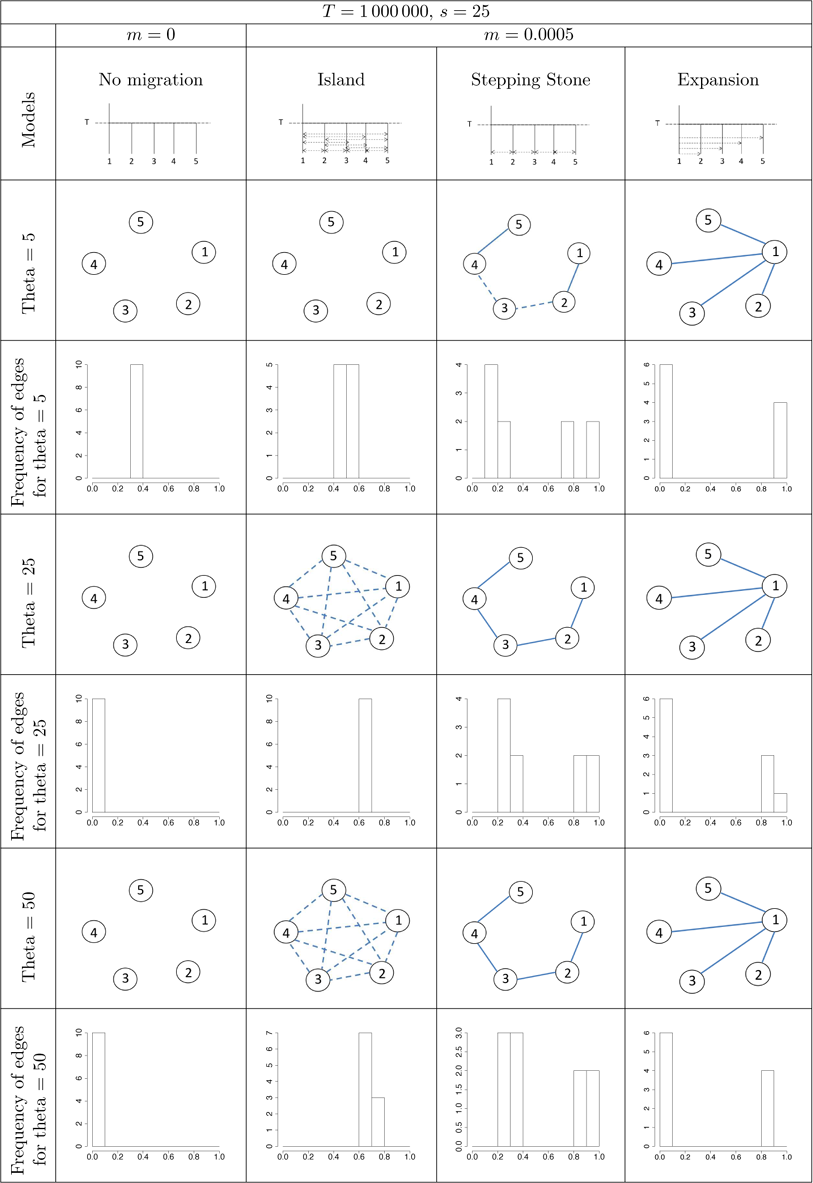
Graphs and histograms resulting of 1000 simulations performed with *T* = 1 000 000, *m* = 0.0005, *s* = 25 and various combinations of models and θ. In the graphs, solid lines correspond to detection probabilities higher than 0.8. Dashed lines correspond to detection probabilities lying from 0.6 to 0.8. The number of solid and dashed lines can be found in the histograms.

**Figure 2:**
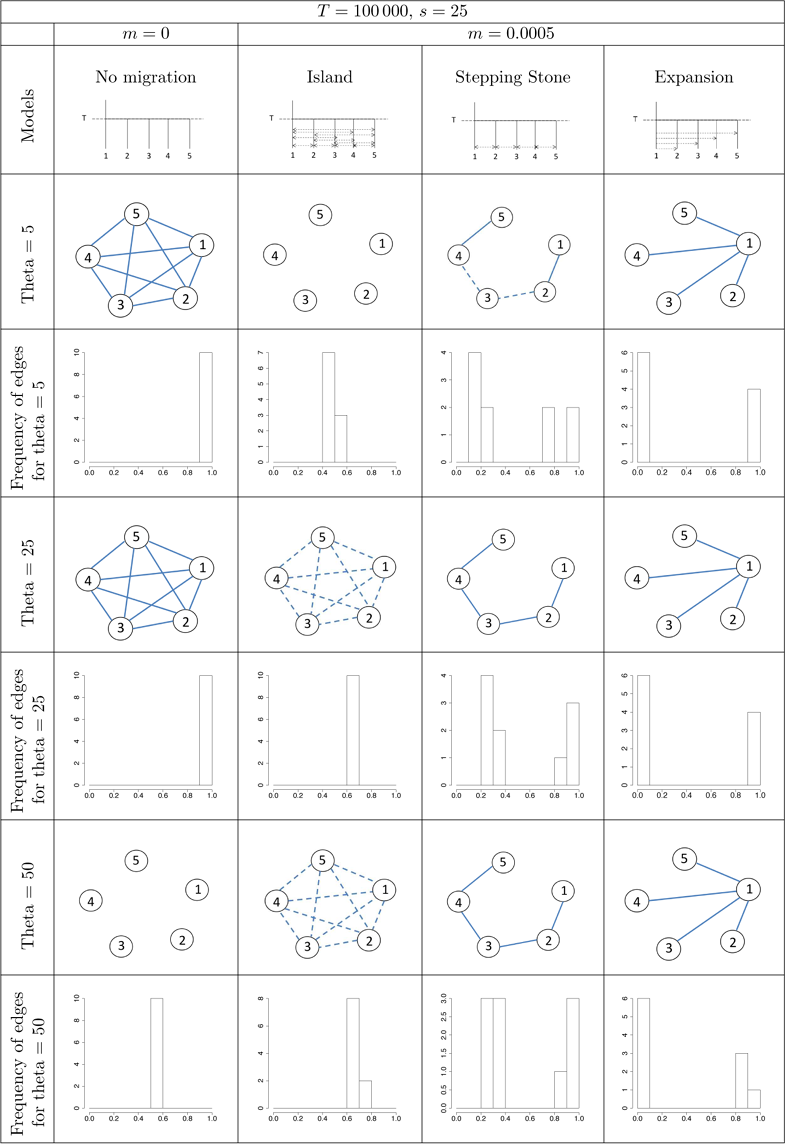
Graphs and histograms resulting of 1000 simulations performed with *T* = 100 000, *m* = 0.0005, *s* = 25 and various combinations of models and θ. In the graphs, solid lines correspond to detection probabilities higher than 0.8. Dashed lines correspond to detection probabilities lying from 0.6 to 0.8. The number of solid and dashed lines can be found in the histograms.

For the model with `no migration` but with a single splitting time, the estimated graphs differed according to *T* and θ. For a high value of *T* (Fig. 1), no edge was detected whatever θ. For a low value of *T* (Fig. 2), the graph was complete (i.e. all nodes are connected) for θ = 5 and θ = 25, while no edge was detected for θ = 50.

Results for the `multi-T` model are presented on Figure 3. With low T, graphs were similar whatever the value of θ. With high *T*, edges 2-3 then 3-4 and 3-5 disappeared when θ increased.

**Figure 3:**
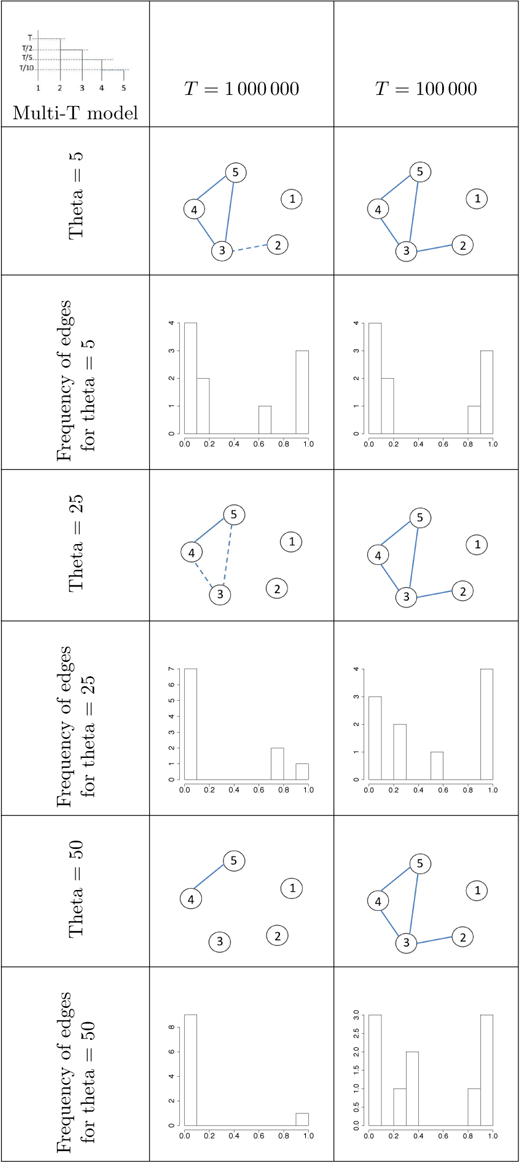
Results of 1000 simulations performed with the “multi-T” model for various combinations of T, θ and *s*. In the graphs, solid lines correspond to detection probabilities higher than 0.8. Dashed lines correspond to detection probabilities lying from 0.6 to 0.8.

## 5 Discussion

The method developed here, was performed on simulated datasets in order to assess its performances. This method was primarily developed for Barcode data which targets a single gene. Nevertheless, this method can be used with a data set composed of several genes.

When *T* = 1 000 000 (Fig. 1) the inferred links recovered the underlying model quite well: no link for the `no migration` model, a path for the `linear Stepping Stone` model, a complete graph for the Island model except for θ = 5, four links from the population 1 for the expansion model and links between recently split populations for the multi-T model. This last configuration highlighted the influence of the splitting time on the results. For *T* decreasing down to 100 000 (Fig. 2), a complete graph was inferred with the `no migration` model as long as θ ≤ 25. Besides, we observed that no link was detected with the `Island` model for θ = 5. This suggests that among-population migrations modified the prior pattern structure and that the new structure is revealed with increasing splitting time *T* or mutation rate θ. Histograms concerning the Island model showed that the detection probabilities of links increased with θ and were close to 0.8 for θ = 50.

Considering the `linear Stepping Stone` model for both values of *T*, the detection probabilities of the two links 2 − 3 and 3 − 4 were very close to 0.8 for the lowest value of θ. This pattern reflected a slight decrease in power of the method when the node degree increased (node degree: number of links linked to this node). Note that this phenomenon was previously described in Villers et al. [2008]. In this case, each node 1 and 5 have one potential link making them easier to detect than the two potential links of nodes 2, 3 and 4.

In our method, we use the SNPs to avoid as much as possible, correlations between the sites. However edges are set between populations if their correlation structure is similar. So assuming independence across sites is not the real assumption on which the analysis relies. Each population is associated to a probability distribution *Q*_*k*_ on [0, 1]^4*L*^ which contains the probability of observing *A, C, G, T* at each polymorphic site. Finding links between two populations given all the others, means that the two random variables *X_*k*_, X_*k*′_* with distribution *Q_*k*_, Q_*k*′_* interact in some way.

Our graph based method takes simultaneously into account the genetic covariance of all populations. Thus, it provides a single heuristic approach to address a wide range of questions in population genetics. Being able to visually inspect the relationships among all populations simultaneously, may prove to be immensely important for exploratory data analysis, data interpretation and the development of subsequent hypothesis.

To conclude, the method presented here completes the range of tools already available in population genetics. Using DNA sequence data, this method which is not based on genetic models brings a new point of view on structured populations. The results were always consistent with biological patterns. However we must keep in mind that a graph is related to a data set and adding nodes or gene sequences will most likely modify the graph structure. Anyway, when DNA sequences are available, our method should be helpful for understanding geographical extension of species, gene flows, biological invasions, and, in epidemiology, to investigate dispersal routes of infectious strains.

## References

T. W. Anderson. An Introduction to Multivariate Statistical Analysis. Wiley-Interscience; 3 edition, 2003.

Yoav Benjamini and Yosef Hochberg. Controlling the False Discovery Rate: A Practical and Powerful Approach to Multiple Testing. Journal of the Royal Statistical Society. Series B (Methodological), 57(1):pp. 289–300, 1995.

C. E Bonferroni. Teoria statistica delle classi e calcolo delle probabilita. In Universita di Firenza, editor, Volume in Onore di Ricarrdo dalla Volta, pages 1–62. 1937.

L L Cavalli-Sforza and A W Edwards. Phylogenetic analysis. Models and estimation procedures. American journal of human genetics, 19(3 Pt 1):233–57, May 1967. ISSN 0002-9297.

R. J. Dyer and J. D. Nason. Population graphs: the graph theoretic shape of genetic structure. Molecular Ecology, 13(7):1713–1727, 2004.

R. A. Fisher. The distribution of the partial correlation coe cient. Metron, 3(3-4):329–332, 1924.

J C Gower. Some distance properties of latent root and vector methods used in multivariate analysis. Biometrika, 53:325–338, 1966.

Gilles Guillot. On the inference of spatial structure from population genetics data. Bioinformatics, 25(14):1796–1801, 2009.

J. Hey. Isolation with migration models for more than two populations. Mol Biol Evol, 27(4): 905–920, 2010.

T Jombart, S Devillard, A-B Dufour, and D Pontier. Revealing cryptic spatial patterns in genetic variability by a new multivariate method. Heredity, 101(1):92–103, July 2008. ISSN 1365-2540. doi: 10.1038/hdy.2008.34. URL http://dx.doi.org/10.1038/hdy.2008.34

M Kimura and G H Weiss. The Stepping Stone Model of Population Structure and the Decrease of Genetic Correlation with Distance. Genetics, 49(4):561–76, April 1964.

J. F. C. Kingman. The coalescent. Stochast Proc App, 13:235–248, 1982.

Edward L Korn, James F Troendle, Lisa M McShane, and Richard Simon. Controlling the number of false discoveries: application to high-dimensional genomic data. Journal of Statistical Planning and Inference, 124(2):379–398, 2004.

Ste↵en L. Lauritzen. Graphical Models, volume 17 of Oxford Statistical Science Series. 1996.

G. Laval and L. Excoer. Simcoal 2.0: a program to simulate genomic diversity over large recombining regions in a subdivided population with a complex history. Bioinformatics, 20(15):2485–2487, 2004.

Alejandro F. Rozenfeld, Sophie Arnaud-Haond, Emilio Hernandez-Garcia, Victor M. Eguiluz, Ester A. Serrao, and Carlos M. Duarte. Network analysis identifies weak and strong links in a metapopulation system. Proceedings of the National Academy of Sciences of the United States of America, 105(48):18824–18829, 2008.

Fanny Villers, Brigitte Schae↵er, Caroline Bertin, and Sylvie Huet. Assessing the validity domains of graphical gaussian models in order to infer relationships among components of complex biological systems. Statistical Applications in Genetics and Molecular Biology, 7(2): 1544–6115, 2008.

Joe Whittaker. Graphical models in applied multivariate statistics. Wiley, 1990. ISBN 0471917508.

S Wright. Evolution in Mendelian Populations. Genetics, 16(2):97–159, March 1931.

